# *Ganoderma formosanum* polysaccharides enhance antitumor immune responses by downregulating the differentiation of myeloid-derived suppressor cells and tumor-associated macrophages

**DOI:** 10.1101/2024.12.23.630057

**Authors:** Jhe-Yu Yang, Kuei-Liang Chen, Yang-Chia Lin, Hsien-Chan Chiu, Hsin-Tien Hsieh, Chun-Jen Chen

**Affiliations:** Department of Biochemical Science and Technology, National Taiwan University, Taipei 10617, Taiwan (R.O.C.); Graduate Institute of Molecular and Cellular Biology, National Taiwan University, Taipei 10617, Taiwan (R.O.C.)

## Abstract

*Ganoderma formosanum* is a native species of *Ganoderma* isolated in Taiwan, and our previous studies showed that a polysaccharide fraction, PS-F2, purified from the submerged culture fluid of *G. formosanum* ATCC 76538 exhibited immunostimulatory and antitumor property. In the current study, we investigated the immunomodulatory and antitumor effects of PS-F2 from a UV-mutated *G. formosanum* variant NTU-1, which produced higher yields of PS-F2 than the original strain. Oral administration of PS-F2 effectively suppressed the growth of colon 26 (CT-26) carcinoma and splenomegaly in tumor-bearing mice without adversely affecting the animals’ health. We found that PS-F2 treatment resulted in augmented cytotoxic T lymphocyte (CTL) while significantly reducing the accumulation of polymorphonuclear myeloid-derived suppressor cells (PMN-MDSCs) and regulatory T (Treg) cells in the spleen. In the tumor, PS-F2 treatment markedly enhanced CTL and Th1 responses, whereas it reduced the accumulation of tumor-associated macrophages (TAMs). Collectively, our data demonstrate that oral treatment of PS-F2 from *G. formosanum* NTU-1 in CT26 tumor-bearing mice activates antitumor immune responses and reduces the accumulation of immunosuppressive cells in the spleen and the tumor, leading to delayed tumor progression.

## Introduction

Increasing numbers of natural products have been shown to function as biological response modifiers (BRMs), particularly in cancer immunotherapy. Compelling data demonstrate that natural products, such as polysaccharide extracts, polyphenols, terpenoids, and cardiotonic steroids, can modulate the immune responses in the host and are promising candidates for cancer therapy ^1^. Natural polysaccharides with bioactive properties have been identified in different organisms, including bacteria, fungi, higher plants, animals, lichens, and algae ^2^. Among them, fungal polysaccharides have attracted great attention, primarily on their immunomodulatory and antitumor effects ^3^. Most fungal polysaccharides with bioactive functions have β-1,3-glucan backbones and may have different sugar compositions, types of linkage, and structures ^4^. The immunostimulatory and antitumor activities of fungal polysaccharides are primarily based on their recognition by host pattern recognition receptors (PRRs) on antigen-presenting cells such as macrophages and dendritic cells, which in turn activate T and B lymphocytes ^5^.

The immune cells within the tumor microenvironment (TME) play important roles in regulating tumorigenesis ^6^. In the cancer-immunity cycle, antigens released from cancer cells can be presented by antigen-presenting cells, and then primed and activated T cells recognizing tumor rejection antigens can target and kill cancer cells ^7^. However, various immunoregulatory mechanisms are also present in the TME to suppress T cell activity and counteract the antitumor immune response ^8^. The primary aims of cancer immunotherapy include the activation of antitumor immunity and the alleviation of the immunosuppressive effect of immunoregulatory cells ^9–12^. Several types of tumor-associated myeloid cells that can suppress T cell activation have been characterized, including tumor-associated macrophages (TAMs), tumor-associated neutrophils (TANs), and myeloid-derived suppressor cells (MDSCs). In the TME, tumor-associated myeloid cells have complex interactions with cancer cells, stromal cells, and other immune cells, resulting in the release of a range of immunosuppressive factors that inhibit the proliferation and cytotoxicity of T cells and NK cells ^12^. Lymphocytes with immunoregulatory functions, including regulatory T (Treg) cells and regulatory B (Breg) cells, also frequently accumulate in the TME and counteract the antitumor immunity ^8^. Therefore, therapeutic strategies to deplete or reprogram the immunosuppressive myeloid and/or lymphoid cells may enhance antitumor responses in patients.

We previously reported that PS-F2, a purified polysaccharide fraction from the submerged culture of *Ganoderma formosanum*, exhibits immunostimulatory and adjuvant activities, promoting a T helper a (Th1)-polarized adaptive immune response in mice ^13–16^. When administered intraperitoneally to tumor-bearing mice, PS-F2 activates tumor-specific cellular and humoral immune responses ^17^. Herein, we show that a fast-growing *G. formosanum* variant NTU-1 was generated, and oral administration of PS-F2 produced by *G. formosanum* NTU-1 also suppressed the growth of CT26 colorectal carcinoma cells in mice. The antitumor response was associated with the activation of Th1 cells and cytotoxic T lymphocytes (CTLs), as well as the reduction in TAMs, MDSCs, and Treg cells.

## Results

### Production and characterization of extracellular polysaccharides from *G. formosanum* NTU-1

In this study, we first aimed to generate fast-growing mutants of *G. formosanum* 76538 by UV mutagenesis. After UV irradiation of the protoplasts, one of the fastest-growing colonies was selected and named *G. formosanum* NTU-1. In the submerged mycelial culture, the crude polysaccharides produced by the NTU-1 variant could be separated into 3 major fractions PS-F1, PS-F2, and PS-F3 (Figure 1A), which resemble the polysaccharides produced by the wild-type strain ^16^. Compared with the wild-type strain, the culture of the variant produced significantly higher levels of crude polysaccharides and PS-F2 (Figure 1B). The major constituents of PS-F2 produced by the NTU-1 variant were D-mannose (46.2 ± 6.3%), D-galactose (36.7 ± 4.1%) and D-glucose (6.2 ± 1.2%), similar to the monosaccharide compositions of PS-F2 from the wild-type strain ^16^. These results indicate that the polysaccharides produced by wild-type and NTU-1 strains exhibit similar properties; however, the NTU-1 variant grows faster and produces more extracellular polysaccharides than the wild-type strain. We then used the PS-F2 polysaccharides produced by *G. formosanum* NTU-1 for the following studies.

**Fig. 1.**
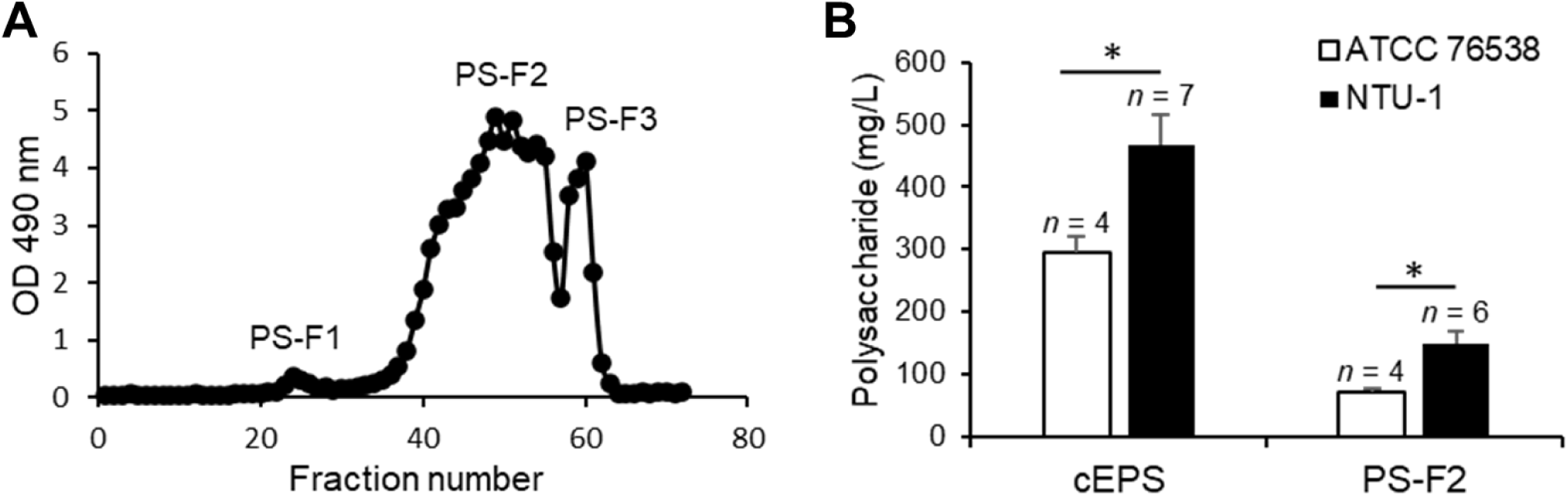
Production and purification of polysaccharides from *G. formosanum* NTU-1. (A) Gel filtration chromatography profile of the extracellular polysaccharides produced in the submerged culture of *G. formosanum* NTU-1. (B) Comparison of the yield of crude extracellular polysaccharides (cEPS) and PS-F2 from *G. formosanum* ATCC 76538 and *G. formosanum* NTU-1.

### Antitumor activity of PS-F2 produced by *G. formosanum* NTU-1

To investigate whether oral administration of PS-F2 could suppress tumor growth in animals, we used the CT26 colorectal carcinoma model in BALB/c mice. Mice were administered p.o. with different doses of PS-F2 (50 mg/kg, 75 mg/kg, and 100 mg/kg) or PBS every other day from day 1 to day 19 after s.c. inoculation of the CT26 tumor. As shown in Figure 2A, starting from day 11 after the tumor challenge, tumor growth was significantly suppressed in mice administered with PS-F2 at 75 mg/kg and 100 mg/kg compared with the PBS group. On day 21 after tumor inoculation, the mice were sacrificed, and the tumors and spleens were completely resected and weighed. The average tumor weights in mice treated with PS-F2 at 75 mg/kg and 100 mg/kg were reduced by 70% and 81% compared with that in PBS-treated mice, respectively (Figure 2B). In addition, PS-F2 treatment at 75 mg/kg and 100 mg/kg also significantly reduced spleen weights in tumor-bearing mice (Figure 2C). Administration of PS-F2 at different doses had no significant effect on mouse body weight (data not shown), indicating that PS-F2 suppressed tumor growth without causing overall toxicity to the animals. These results demonstrated that oral administration of PS-F2 effectively reduced splenomegaly and delayed tumor growth in mice bearing the CT26 tumor.

**Fig 2.**
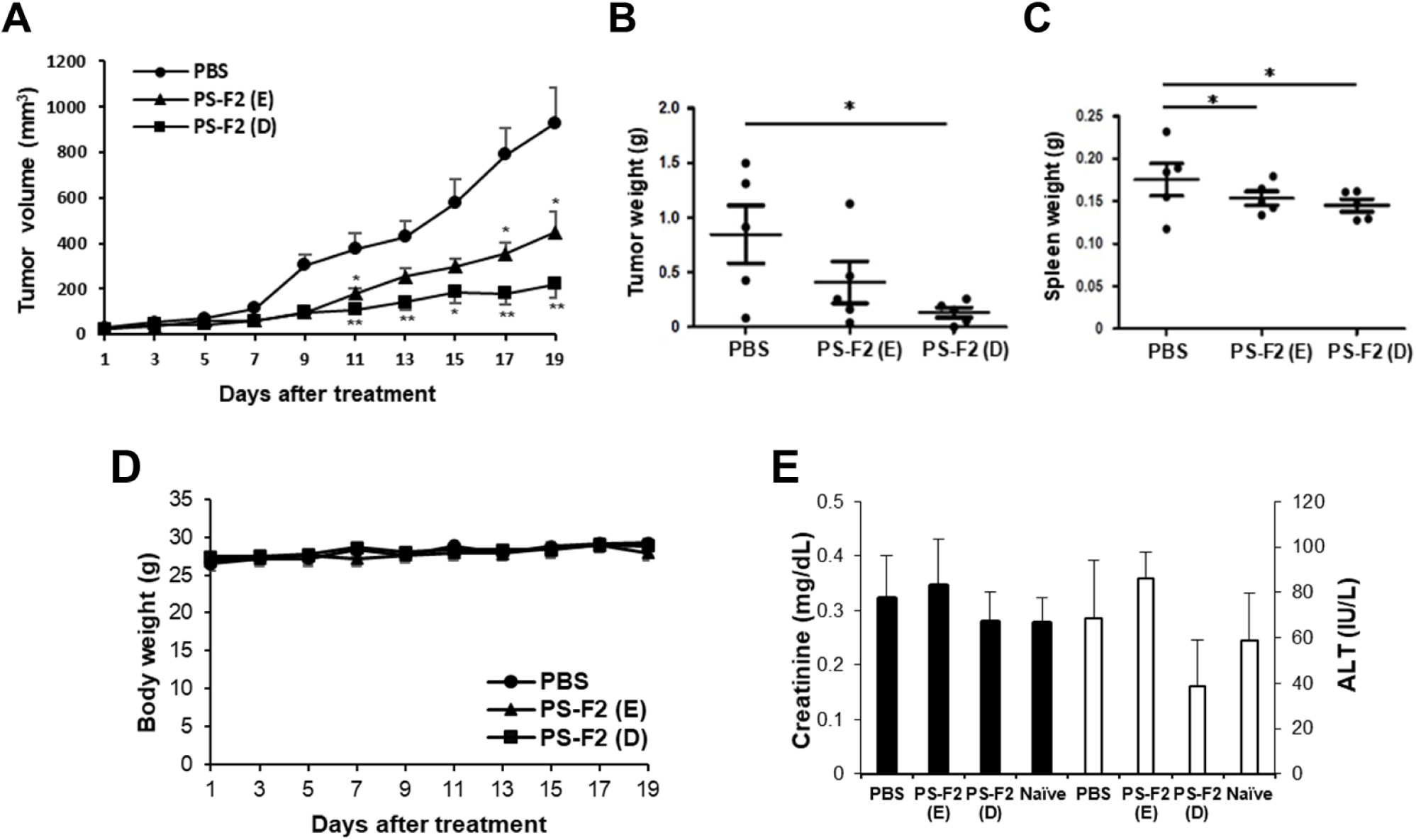
Antitumor effect of PS-F2 in CT26 tumor-bearing mice. CT26 tumor cells were inoculated s.c. into BALB/c mice on day 0. (A-C) Mice were administered p.o. with PS-F2 (50, 75, or 100 mg/kg) or PBS every other day from day 1 to day 19, and average tumor volume was monitored (A). On day 21, the mice were sacrificed, and the tumors (B) and spleens (C) were resected and weighed (*n* = 4 or 5). \**P* < 0.05, \*\**P* < 0.01.

### PS-F2 administration suppresses the growth of established tumors in mice

To investigate whether PS-F2 exhibited an antitumor effect when given to mice with established CT26 tumors, mice were inoculated s.c. with CT26 tumor cells, and when the average tumor size reached 5 mm in diameter, mice were administered p.o. with PBS or PS-F2 (75 mg/kg) daily or every other day for 3 weeks. As shown in Figure 3A, from day 11 after treatment, tumor growth was significantly suppressed in PS-F2-treated animals compared with the PBS group, and the suppressive effect was more evident in mice receiving daily administration of PS-F2. On day 21 after treatment, the mice were sacrificed, and the tumors and spleens were completely resected and weighed. Compared with the PBS group, the average tumor weights in PS-F2-treated groups were reduced by 52% and 84% in mice receiving PS-F2 treatment every other day or daily, respectively (Figure 3B). Administration of PS-F2 also resulted in reduced spleen weights in CT26 tumor-bearing mice (Figure 3C). In contrast to its suppressive effect on tumor growth, oral administration of PS-F2 had no significant effect on mouse body weight (Figure 3D), and serum levels of creatinine and ALT activities were in the normal range (Figure 3E). These results indicated that PS-F2 administration suppressed tumor growth without causing general toxicity to the host.

**Fig 3.**
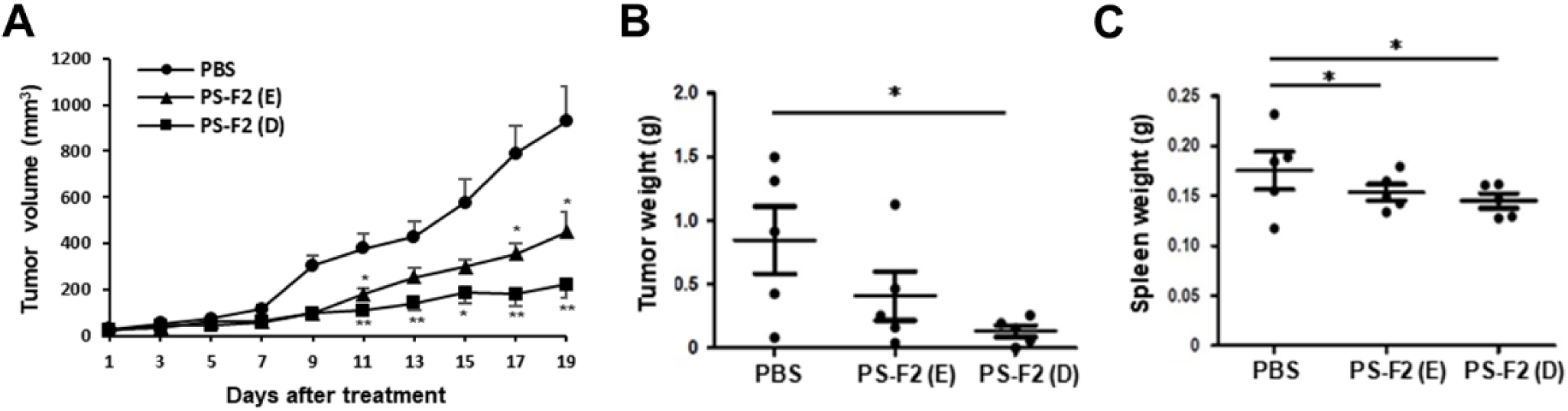
Antitumor effect of PS-F2 in mice bearing established CT26 tumors. CT26 tumor cells were inoculated s.c. into BALB/c mice, and when the average tumor size reached 5 mm in diameter, mice were orally administered with PBS or PS-F2 (75 mg/kg) by gavage every other day [PS-F2 (E)] or daily [PS-F2 (D)]. (A) The average tumor volume was measured every other day. (B, C) On day 19, the mice were sacrificed, and the tumors (B) and spleens (C) were resected and weighed (*n* = 5). The data shown are representative of 3 independent experiments. \**P* < 0.05, \*\**P* < 0.01.

### PS-F2 administration induces the activation of Th1 cells and CTLs in tumor-bearing mice

To investigate whether PS-F2 administration exerts the antitumor effect via inducing a Th1-polarized immune response and CTL activation, we analyzed the CD4^+^IFN-γ^+^ Th1 cells and CD8^+^IFN-γ^+^ CTLs in spleens and tumors by flow cytometry. As shown in Figure 4A, PS-F2 administration resulted in an increase in the frequency of IFN-γ-producing CD4^+^ cells in splenocytes and tumor-infiltrating lymphocytes (TILs). PS-F2 treatment also augmented the population of CD8^+^IFN-γ^+^ CTLs in spleens and tumors (Figure 4B). The enhanced Th1 and CTL responses were more evident when mice received PS-F2 treatment daily versus every other day, indicating that dosing PS-F2 more frequently could promote a stronger activation of Th1 cells and CTLs, resulting in a greater antitumor effect in mice.

**Fig 4.**
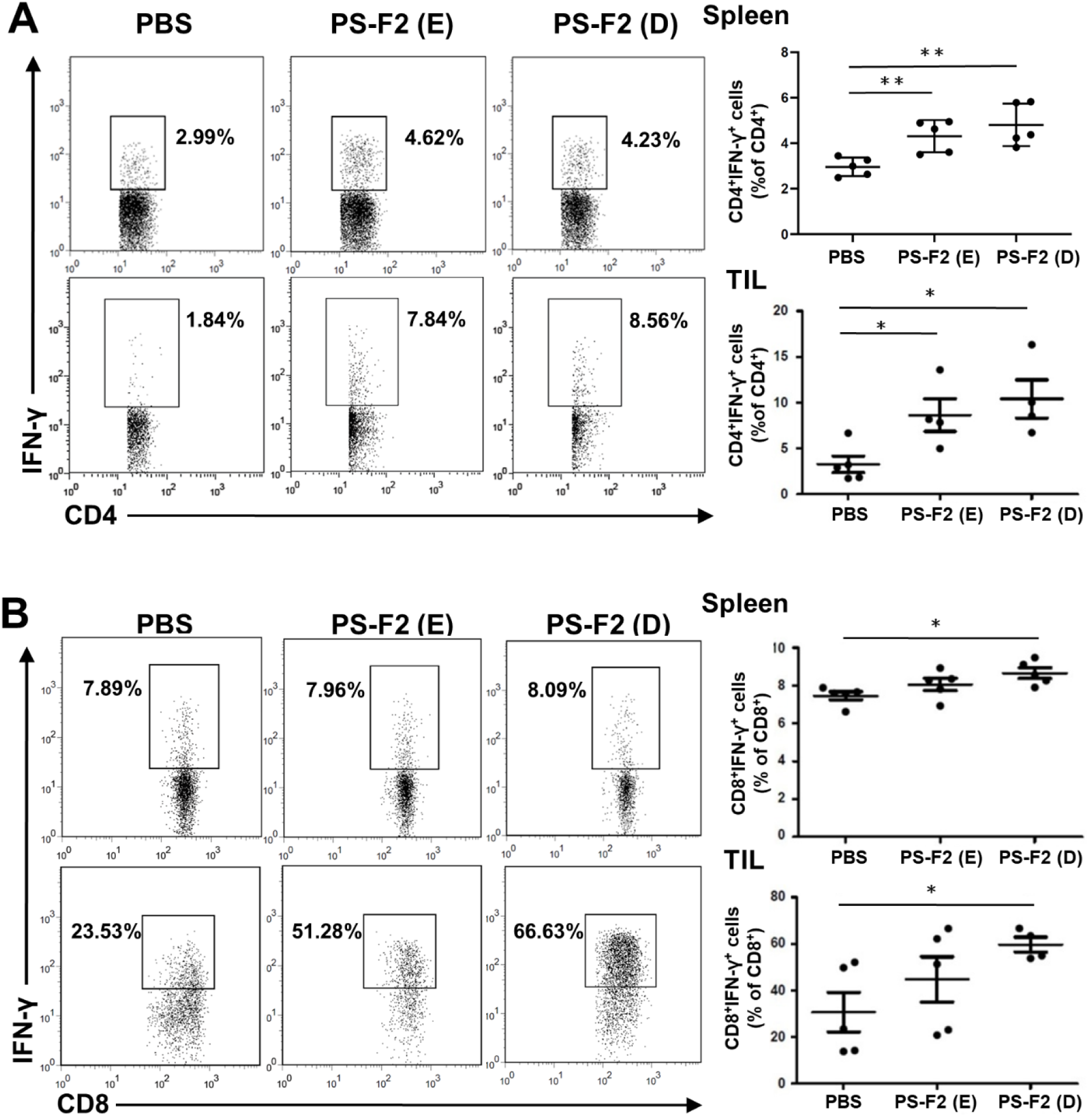
Therapeutic administration of PS-F2 enhanced Th1 and CTL responses in CT26 tumor-bearing mice. CT26 tumor cells were inoculated s.c. into BALB/c mice, and when the average tumor size reached 5 mm in diameter, mice were administered p.o. with PBS or PS-F2 (75 mg/kg) every other day [PS-F2 (E)] or daily [PS-F2 (D)] from day 1 to day 19. On day 19, mice were sacrificed, and single-cell suspensions from spleens and tumor tissues were stained with corresponding antibodies to evaluate the proportions of CD4^+^IFN-γ^+^ Th1 cells (A) and CD8^+^IFN-γ^+^ CTLs (B) in splenocytes and tumor-infiltrating lymphocytes (TIL) by flow cytometry. Cells were gated on CD4^+^ (A) or CD8^+^ (B) cells. Data shown are representative of 3 independent experiments. \**P* < 0.05, \*\**P* < 0.01.

### PS-F2 treatment alters the proportions of MDSCs and TAMs in tumor-bearing mice

Having observed that PS-F2 treatment could stimulate the activation of Th1 and CTL responses, we next investigated whether this effect was associated with altering cell populations with immunosuppressive functions. As shown in Figure 5a, PS-F2 administration resulted in a significant decrease in the proportion of CD11b^+^Ly6G^+^Ly6C^int^ PMN-MDSCs in the spleen. The frequency of PMN-MDSCs in the tumor was also moderately reduced when mice received daily treatment of PS-F2 (Figure 5B). In addition, PS-F2 administration also markedly reduced the proportion of CD11b^hi^F4/80^+^ TAMs in the tumor (Figure 5C). In contrast, PS-F2 treatment did not affect the proportion of CD11b^+^Ly-6G^−^Ly6C^hi^ M-MDSCs in the spleen (Figure 5A) and tumor (Figure 5B). These data indicate that PS-F2 could suppress the accumulation of myeloid-derived immunosuppressive cells, notably PMN-MDSCs and TAMs, in tumor-bearing mice.

**Fig 5.**
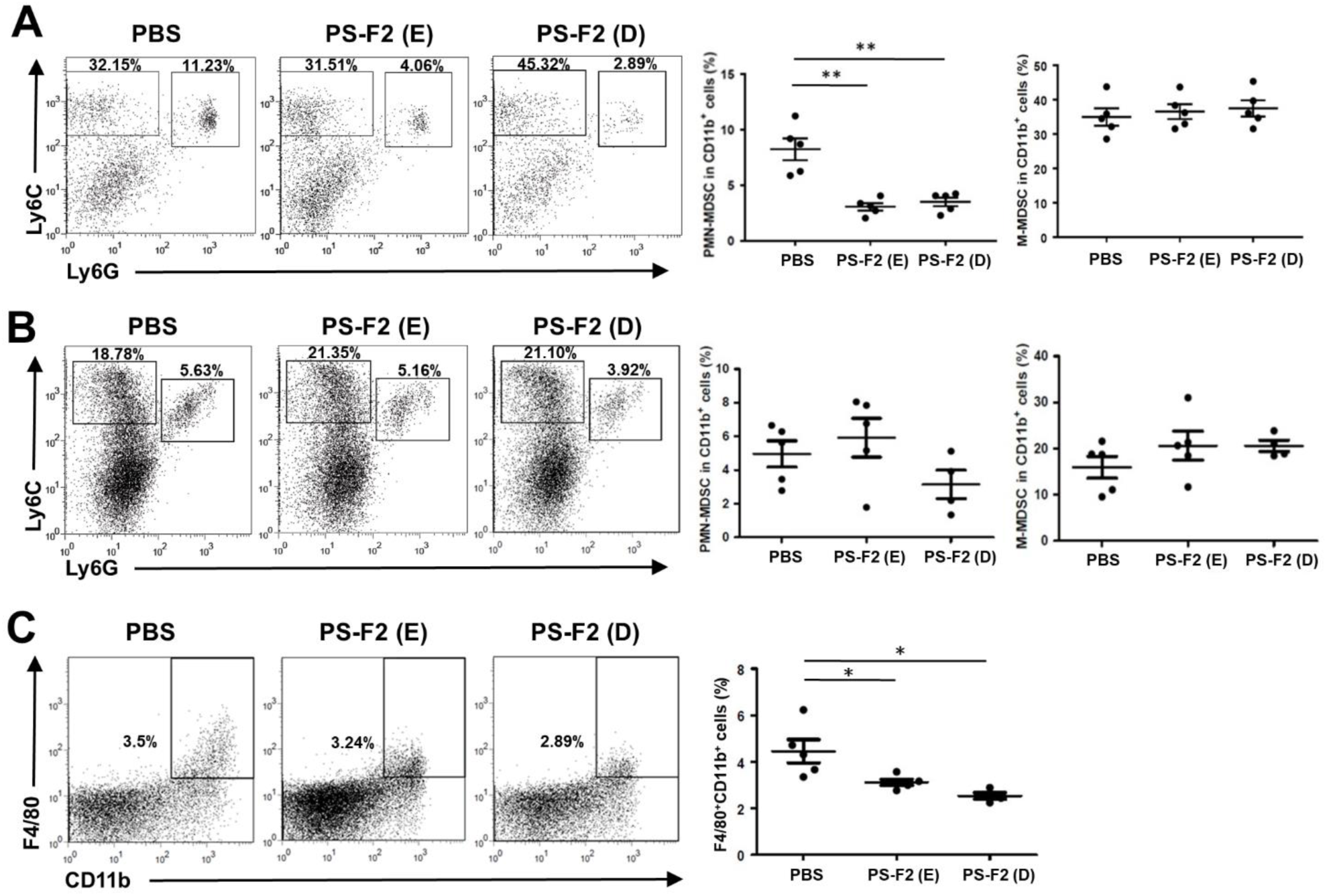
Therapeutic administration of PS-F2 downregulates the differentiation of PMN-MDSCs and TAMs in CT26 tumor-bearing mice. CT26 tumor cells were inoculated s.c. into BALB/c mice, and when the average tumor size reached 5 mm in diameter, mice were administered p.o. with PBS or PS-F2 (75 mg/kg) every other day [PS-F2 (E)] or daily [PS-F2 (D)] from day 1 to day 19. On day 19, mice were sacrificed, and single cell suspensions from spleens and tumor tissues were analyzed by flow cytometry. (A, B) Splenocytes (A) and tumor tissue cells (B) were analyzed for PMN-MDSCs (CD11b^+^Ly6G^+^Ly6C^int^) and M-MDSCs (CD11b^+^Ly-6G^−^ Ly6C^hi^). Cells were gated on CD11b^+^ cells. (C) Tumor tissue cells were analyzed for TAMs (CD11b^hi^F4/80^+^). Data shown are representative of 2 independent experiments. \**P* < 0.05, \*\**P* < 0.01.

### PS-F2 treatment impacts the differentiation of Treg cells in tumor-bearing mice

Treg cells are frequently found to increase in tumor-bearing animals and suppress antitumor immunity. We found that splenic CD4^+^CD25^+^Foxp3^+^ Treg cells were markedly increased in tumor-bearing mice compared with naïve animals (data not shown), and PS-F2 administration could reduce the frequency of splenic Treg cells in a dose-dependent manner (Figure 6A). However, PS-F2 treatment had less effect on the proportion of Treg cells in the tumor tissue. Consistent with the lower numbers of Treg cells in the spleen, PS-F2-treated animals also had lower levels of TGF-β in the serum (Figure 6B). Taken together, these data suggest that in tumor-bearing mice, oral administration of PS-F2 suppressed the differentiation of cells with immunosuppressive properties, including MDSCs, TAMs, and Treg cells, while augmenting CTL and Th1 responses, thus leading to the attenuation of tumor growth.

**Fig 6.**
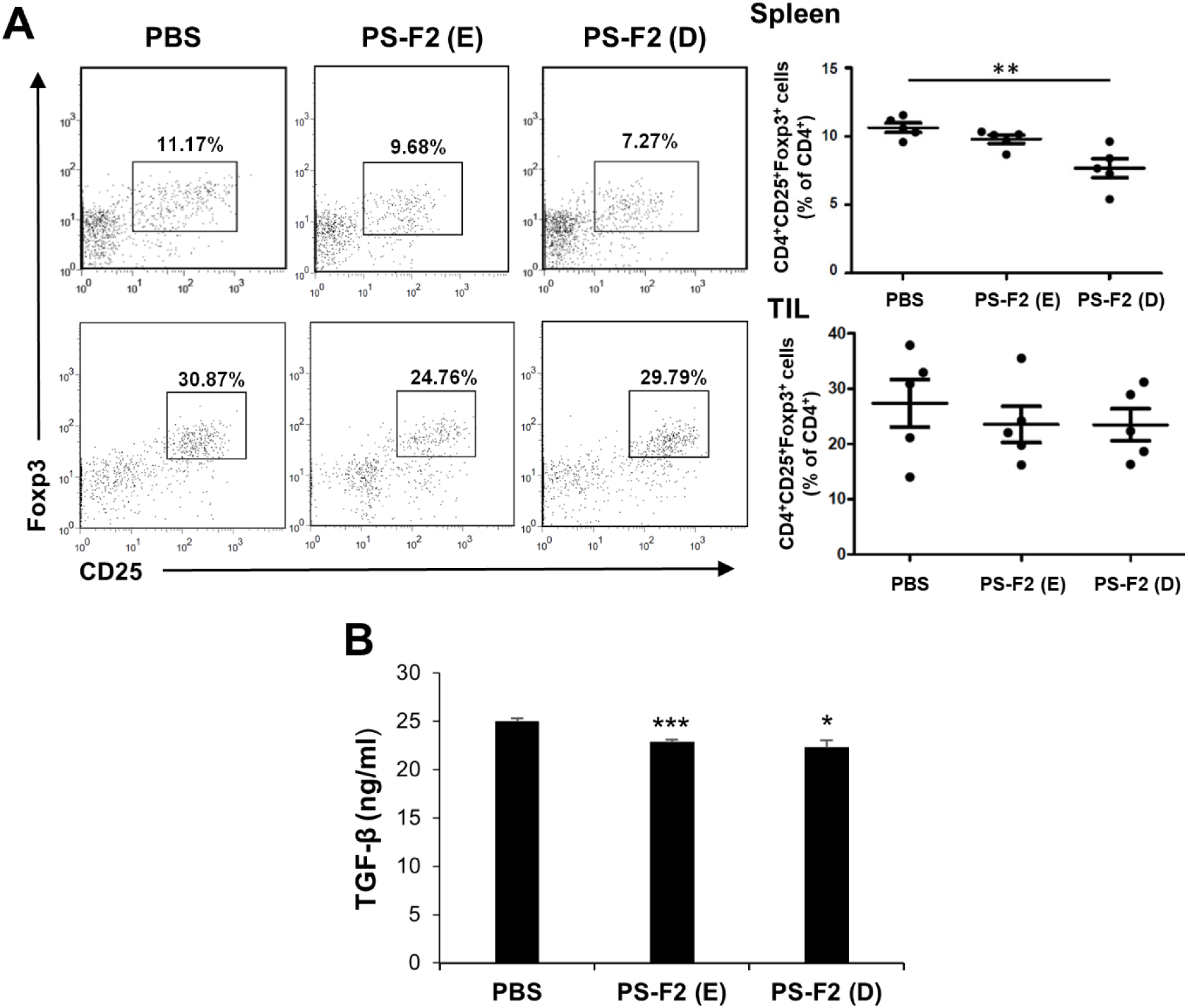
PS-F2 administration impacts the differentiation of Treg cells and the level of serum TGF-β in CT26 tumor-bearing mice. CT26 tumor cells were inoculated s.c. into BALB/c mice, and when the average tumor size reached 5 mm in diameter, mice were administered p.o. with PBS or PS-F2 (75 mg/kg) every other day [PS-F2 (E)] or daily [PS-F2 (D)] from day 1 to day 19. (A) On day 19, mice were sacrificed, and single-cell suspensions from spleens and tumor tissues were stained with corresponding antibodies to evaluate the proportions of Treg (CD4^+^CD25^+^Foxp3^+^) cells in splenocytes and tumor-infiltrating lymphocytes (TIL). Cells were gated on CD4^+^ cells. (B) On day 19, serum levels of TGF-β were analyzed by ELISA. The data shown are representative of 2 independent experiments. \**P* < 0.05, \*\**P* < 0.01, \*\*\**P* < 0.001.

## Discussion

We have previously reported that PS-F2, a polysaccharide fraction purified from the submerged culture broth of *G. formosanum*, could activate antitumor immune responses when continuously injected into tumor-bearing mice ^17^. In this study, we first generated *G. formosanum* NTU-1, a new variant that produced more polysaccharides than the wild-type strain, and we further investigated the antitumor effect when PS-F2 was orally administered. Our results showed that PS-F2 suppressed tumor growth and splenomegaly when given to mice with established CT26 tumors. PS-F2 treatment induced the activation of Th1 cells and CTLs in the spleen and tumor, while significantly reducing the accumulation of PMN-MDSCs, Treg cells, and TAMs in these tissues. Together, these immunomodulatory mechanisms may contribute to the antitumor effect of PS-F2.

The primary aims of cancer immunotherapy include activating antitumor immunity and alleviating the immunosuppressive effect imposed by cancer cells and immunoregulatory cells. Our data showed that administration of PS-F2 to tumor-bearing mice significantly increased the percentage of IFN-γ-producing CD4^+^ and CD8^+^ cells in the spleen and tumor, indicating the activation of tumor-targeting effector T cell responses. In contrast, the accumulation of MDSCs, particularly PMD-MDSCs, decreased in the spleen and tumor. Furthermore, PS-F2 treatment also reduced the accumulation of Treg cells in the spleen and TAMs in the tumor. Although the frequency of Treg cells in the tumor was not significantly altered in PS-F2-treated mice, it remains possible that the immunosuppressive function of Treg cells could be affected. Such an effect was observed when tumor-bearing mice were treated with yeast β-glucan ^18^. A recent study showed that the crude extracellular polysaccharides from *G. formosanum* (GF-EPS) exerted antitumor activity in mice bearing Lewis lung carcinoma (LLC), which was associated with increased splenic NK cell population, while the frequencies of CD4^+^ and CD8^+^ T cells were not affected ^19^. It is possible that the crude GF-EPS and purified PS-F2 have different effects on the host immune system, however, since the former study did not carefully analyze the activation status of T cells, it remains possible that T cells also contribute to the antitumor effect against LLC. Similar to our observation, the administration of Ganoderma lucidum polysaccharide (GLP) to tumor-bearing mice also resulted in an increase in CTL and Th1 cells and a decrease in Treg cells in the spleen and tumor ^20^.

Our previous studies showed that when PS-F2 was used as an adjuvant and given to animals via intraperitoneal immunization, PS-F2 promoted the generation of antigen-specific Th1 and CTL responses ^14^. This adjuvant property of PS-F2 may also stimulate antitumor immune responses when PS-F2 is continuously injected into tumor-bearing mice ^17^. In this study, our results showed that PS-F2 exhibited antitumor and immunomodulatory activities when delivered via the gastrointestinal route. In the gastrointestinal tract, polysaccharides with β linkages such as PS-F2 are likely utilized by gut microbiota; therefore, we speculate that PS-F2 may exert different biological effects when delivered systemically or orally into the host. Gut microbiota plays an important role in modulating the efficacy of various antitumor agents ^21^. Recent studies have indicated that the antitumor effect of orally ingested polysaccharides, including the polysaccharides from *G. lucidum*, was in part due to the modification of gut microbiota ^20,22^. Whether orally administered PS-F2 alters the composition of gut microbiota and how it may contribute to the antitumor effect of PS-F2 remains to be explored.

Various fungal polysaccharides, primarily from yeasts and mushrooms, have been shown to exhibit potent immunomodulatory and antitumor activities ^5^. Different fungal polysaccharides have diverse structural characteristics, mainly involving monosaccharide composition, degree of branching, molecular weight, and conformation. Polysaccharides from *Ganoderma* have been widely studied for their immunomodulatory and antitumor activities, and most studies have investigated the polysaccharides extracted by hot water from the fruiting bodies of *G. lucidum*, *G. tsugae*, and *G. atrum* ^23^. PS-F2 produced and purified from the submerged culture of *G. formosanum* is a heteroglycan mainly composed of D-mannose, D-galactose, and D-glucose ^16^. PS-F2 differs from the polysaccharides produced by other *Ganoderma* species (e.g. *G. lucidum*) in which D-glucose is usually the major component ^24^. Production of polysaccharides from the submerged mycelial culture of *Ganoderma* can significantly shorten the culture period compared with the culture of fruiting bodies. Therefore, PS-F2 has its unique advantage for future development as a BRM for cancer immunotherapy.

## Materials and methods

### Cell lines and animals

Murine colon 26 (CT26) cells were maintained in RPMI 1640 supplemented with 10% fetal bovine serum (FBS). Male BALB/c mice (6 weeks old) were purchased from the National Laboratory Animal Center (Taipei, Taiwan). The animal studies were approved by the Institute Animal Care and Use Committee of National Taiwan University, and all mice were kept in the animal facility of the College of Life Science at National Taiwan University.

### Generation of *G. formosanum* NTU-1

*G. formosanum* Chang et Chen (ATCC 76538) was cultured in 2.4% potato dextrose broth (PDB) for 3 days, and the mycelia were harvested and digested with an enzyme solution (1% kitalase/ml, 0.5% driselase, 0.6 M mannitol, pH 6.5) at 30 ^°^C for 2.5 h. After centrifugation at 4000 x g for 5 min, the protoplasts were resuspended in 0.6 M mannitol (1 x 10^6^ cells/ml) and subjected to UV irradiation for 0, 20, 40, and 60 sec. The UV-irradiated protoplasts were spread onto potato dextrose agar (PDA) containing 0.6 M mannitol and incubated at 25 ^°^C for 9 days, and one of the fastest-growing colonies was selected. The variant was named *G. formosanum* NTU-1 and used for the following studies.

### *G. formosanum* NTU-1 polysaccharides and reagents

The submerged mycelial culture of *G. formosanum* NTU-1 was performed in a 5-L jar fermentor containing MEYE medium (5% glucose, 1.5% malt extract, 0.5% yeast extract, pH 4.5) for 7 days. Polysaccharides in the cleared broth were precipitated in 75% ethanol and purified using a Sepharose CL-6B (Cytiva, Marlborough, MA, USA) column as previously described ^16^, and the major polysaccharide fraction PS-F2 was collected. The purified PS-F2 used for in vitro experiments was further passed through the Detoxi-Gel Endotoxin Removing Gel (Thermo Fisher Scientific, Waltham, MA, USA), and the endotoxin level in the samples was determined to be < 0.3 EU/mg by the Pyrotell *Limulus* Amebocytes Lysate (LAL) test (Associates of Cape Cod, Falmouth, MA, USA). Kitalase was purchased from Fujifilm Wako Pure Chemical Corporation (Osaka, Japan). Driselase was purchased from Sigma-Aldrich (St. Louis, MO, USA). DMEM and RPMI 1640 were purchased from HyClone (Logan, UT, USA). FBS was purchased from Biological Industries (Beit-Haemek, Israel). Phosphate-buffered saline (PBS) was purchased from Corning (Corning, NY, USA). All other chemicals were purchased from commercial sources at the highest purity available.

### Tumor challenge and PS-F2 treatment in mice

CT26 cells (5 x 10^5^) in PBS were injected subcutaneously (s.c.) at the right flank of BALB/c mice. To determine the effect of PS-F2 on tumor growth, mice were administered orally (p.o.) with PBS or PS-F2 (50, 75 or 100 mg/kg in PBS) every other day starting from day 1 to day 21 after tumor inoculation. To determine whether PS-F2 administration affects the growth of established tumors when the average tumor size reached 5 mm in diameter, mice were administered p.o with PBS or PS-F2 (75 mg/kg) daily or every other day. Tumor size was measured using a vernier caliper, and tumor volume was calculated using the formula *V*= (length x width^2^)/2. On day 19 after treatment, mice were sacrificed, and tumors and spleens were completely resected and weighed. Splenocytes and single-cell suspension of tumor tissues were harvested to analyze immune cell populations.

### Preparation of single-cell suspension from the tumor tissue

Tumors were excised, minced into small pieces, and immersed in the digestion mixture containing 5% FBS in RPMI 1640, 1 mg/ml collagenase type I (Sigma-Aldrich), and 0.02 mg/ml DNase Ⅰ (Sigma-Aldrich). This mixture was incubated at 37℃ for 30 min on a rotating platform. The digested cells were filtered through 70-μm cell strainers (Corning) and washed twice with ice-cold PBS. The remaining red blood cells were lysed in the ACK hemolysis buffer.

### Flow cytometric analysis

To analyze cell surface markers, single-cell suspensions were first incubated with mAb 2.4G2 for 30 min to block FcγRIIB/III receptors, followed by incubation with various mAbs (all from BioLegend) as indicated for 30 min at 4℃. The following mAbs were used: CD11b-PE, Ly6G-FITC, Ly6C-APC, CD4-FITC, CD25-APC, F4/80-PE and CD8-APC. To analyze intracellular IFN-γ expression, cells were first stimulated with phorbol-12-myristate-13-acetate (PMA, 50 ng/ml), ionomycin (1 μg/ml) and brefeldin A for 4 h, stained with anti-CD4-FITC and anti-CD8-APC, fixed and permeabilized using the BD Cytofix/Cytoperm Fixation/Permeablization Kit (BD Biosciences, San Jose, CA, USA), and stained with anti-IFN-γ-PE. To analyze intracellular Foxp3 expression, cells were stained with anti-CD4-FITC, fixed and permeabilized using the eBioscience Foxp3/Transcription Factor Staining Buffer Set (Thermo Fisher Scientific), and stained with anti-Foxp3-PE (Thermo Fisher Scientific). Samples were analyzed on a FACSCanto Ⅱ flow cytometer (BD Biosciences), and data were analyzed using FlowJo software (BD Biosciences).

### Statistical analysis

Statistical analysis was performed using an unpaired, two-tailed Student’s *t*-test, and *P* < 0.05 was considered statistically significant. Data are reported as mean ± SEM.

### Data availability

The data presented in this study are available on request from the corresponding author.

## Acknowledgments

We thank the NTU TechComm flow cytometry core facility.

## Author contributions

Conceptualization, J.-Y.Y. and C.-J.C.; methodology, J.-Y.Y., K.-L.C., and C.-J.C.; validation, J.-Y.Y. and C.-J.C.; formal analysis, J.-Y.Y. and C.-J.C.; investigation, J.-Y.Y., Y.-C.L., H.-C.C. and C.-J.C.; resources, C.-J.C.; data curation, J.-Y.Y. and C.-J.C.; writing, J.-Y.Y. and C.-J.C.; C.-J.C.; visualization, J.-Y.Y. and H.-T.H.; supervision and funding acquisition, C.-J.C. All authors have read and agreed to the published version of the manuscript.

